# High throughput single-cell RNA sequencing of intact adult cardiomyocytes and non-myocytes using a split-pool approach

**DOI:** 10.64898/2026.04.28.721288

**Authors:** Yang Hu, Rijan Gurung, Stefan Mueller, Erielle Villanueva, Justus Stenzig, Nirmala Rayan, Tuan Danh Anh Luu, Syahfiqah Nur, Brandon Yong Chen Tan, Boxiang Liu, Haojie Yu, Hyungwon Choi, Roger Sik Yin Foo, Matthew Ackers-Johnson

**Author notes:** Senior author (Matthew Ackers-Johnson).

## Abstract

**MOTIVATION:** Adult cardiomyocytes are difficult to profile by whole-cell single-cell RNA sequencing because of their large size and fragility, which make them poorly compatible with standard workflows. Current approaches for adult cardiomyocyte transcriptomics often require a trade-off between data quality and throughput, thus, studies instead rely heavily on sequencing of nuclei alone. Therefore, we set out to develop a high-quality and scalable workflow for adult heart cells using in-cell ligation and split-pool barcoding strategies to address this methodological gap. This workflow may be further generalisable to other large cell types or samples containing cell populations with highly unequal RNA content.

**SUMMARY:** Adult cardiomyocytes are difficult to profile by whole-cell single-cell RNA sequencing (scRNA-seq). Here, we developed a high-quality and scalable workflow for adult heart cells using in-cell ligation and split-pool barcoding. We identified per-cell RNA content as a significant variable that must be accounted for. Separation of cardiomyocytes (large cells) and non-cardiomyocytes (small cells) before library construction, and allocation of deeper sequencing to cardiomyocytes, produced high-quality whole-cell datasets for both compartments. Compared with single-nucleus RNA sequencing, whole-cell cardiomyocyte profiling better recovered metabolic, mitochondrial, cytoplasmic translational, and contractile gene programs. This workflow provides a practical method for scalable, high-quality cardiomyocyte whole-cell scRNA-seq and offers general strategies for other large cell types or samples containing cell populations with highly unequal RNA content.

## INTRODUCTION

Single-cell RNA sequencing (scRNA-seq) has become an important tool for studying cardiac biology(1). Cardiomyocytes (CMs) are central to cardiac physiology and disease. However, sequencing intact adult CMs at single-cell resolution remains technically challenging. Adult CMs are large and fragile, making isolation difficult, and poorly compatible with standard microfluidic workflows (e.g., 10x Chromium). As a result, standard whole-cell scRNA-seq approaches are often difficult to apply to adult CMs.

Accordingly, single-nucleus RNA sequencing (snRNA-seq) has become the most commonly used approach for adult CM transcriptomics. snRNA-seq overcomes the cell-size barrier, can be applied to frozen tissues, and can reduce dissociation-induced stress responses and cell-type capture bias in some settings(2, 3). It is also compatible with multiomic assays(2). However, snRNA-seq has important limitations: it loses cytoplasmic transcripts, typically detects fewer genes per cell than whole-cell methods, is susceptible to transcriptional bursting, and introduces nuclear expression bias(2). In addition, nuclear counts are not equivalent to cell counts in multinucleated cells, which is particularly relevant in species with sizeable multinucleated CM populations, including humans and rodents(2, 4). These features make snRNA-seq powerful but not interchangeable with whole-cell CM scRNA-seq.

Several whole-cell scRNA-seq approaches for adult CMs have been reported, but each has practical limitations. Conventional droplet-based platforms such as 10x Chromium are generally suitable for smaller CMs (e.g., embryonic, neonatal, or iPSC-derived CMs), but not for intact adult mammalian CMs due to cell-size constraints(5). FACS-based approaches have reported adult CM scRNA-seq (for example, FACS sorting followed by SORT-seq library preparation)(6), but the data exhibit signatures of compromised quality associated with degraded cardiomyocyte integrity including high mitochondrial transcript fractions. Large-particle FACS (LP-FACS) followed by mcSCRB-seq improves recovery of intact rod-shaped CMs and produced higher-quality whole-cell CM transcriptomic data, but throughput remains limited(7-9). Nanowell-based platforms such as Takara Bio iCELL8 also demonstrate the feasibility of whole-cell adult CM scRNA-seq, but data quality again appears limited in practice, with high mitochondrial transcript fractions persisting in published datasets(10). By contrast, manual picking workflows can yield very high-quality adult CM transcriptomic data, but are labor-intensive and low-throughput(11, 12). Taken together, current whole-cell CM scRNA-seq methods are often constrained by a choice of either lower data quality or lower throughput.

These limitations underscore the need for a whole-cell CM scRNA-seq method that achieves both high data quality and scalable throughput. Here, we develop a whole-cell scRNA-seq workflow for adult mouse heart cells using in-cell ligation and split-pool barcoding. Per-cell RNA content is identified as a key variable to consider, such that co-processing of CM and non-CM cells otherwise resulted in CM-skewed libraries due to extreme differences in RNA content. In contrast, separation of CM and non-CM cells up to library construction, with subsequent deeper sequencing of CM libraries, enabled high-quality whole-cell CM and non-CM datasets. Benchmarking of our whole-cell CM scRNA-seq data against a snRNA-seq dataset defined method-dependent strengths. This workflow is particularly suitable for studies of CM metabolic, mitochondrial, cytoplasmic, translational, and contractile biology, whereas snRNA-seq remains a powerful option for frozen or difficult to dissociate tissues and for insights into nuclear transcriptional regulation.

## RESULTS

### Co-processing CMs and non-CMs obscures barcode separation and under-represents non-CMs

Parse Evercode^TM^ split-pool barcoding was selected as a commercially available, non-microfluidic technology to investigate applicability to whole-cell CM scRNA-seq. Several modifications were made to manufacturer protocol to optimise for large-cell input, which are detailed in *Methods*. We first attempted to co-process adult mouse ventricular CMs and non-CMs in a single workflow (Figure 1A). Upon sequencing of this mixed library, barcode-rank plots, which aid visualisation of cell-to-background signal discrimination, were observed to lack a distinctive “cliff and knee” configuration that is expected of high-quality sampling (Figure 1B; Table 1). Sequencing saturation was low, and saturation curves did not approach a plateau (Figures 1C and 1D; Table 1), suggesting insufficient coverage for the complexity of the mixed library.

**Figure 1.**
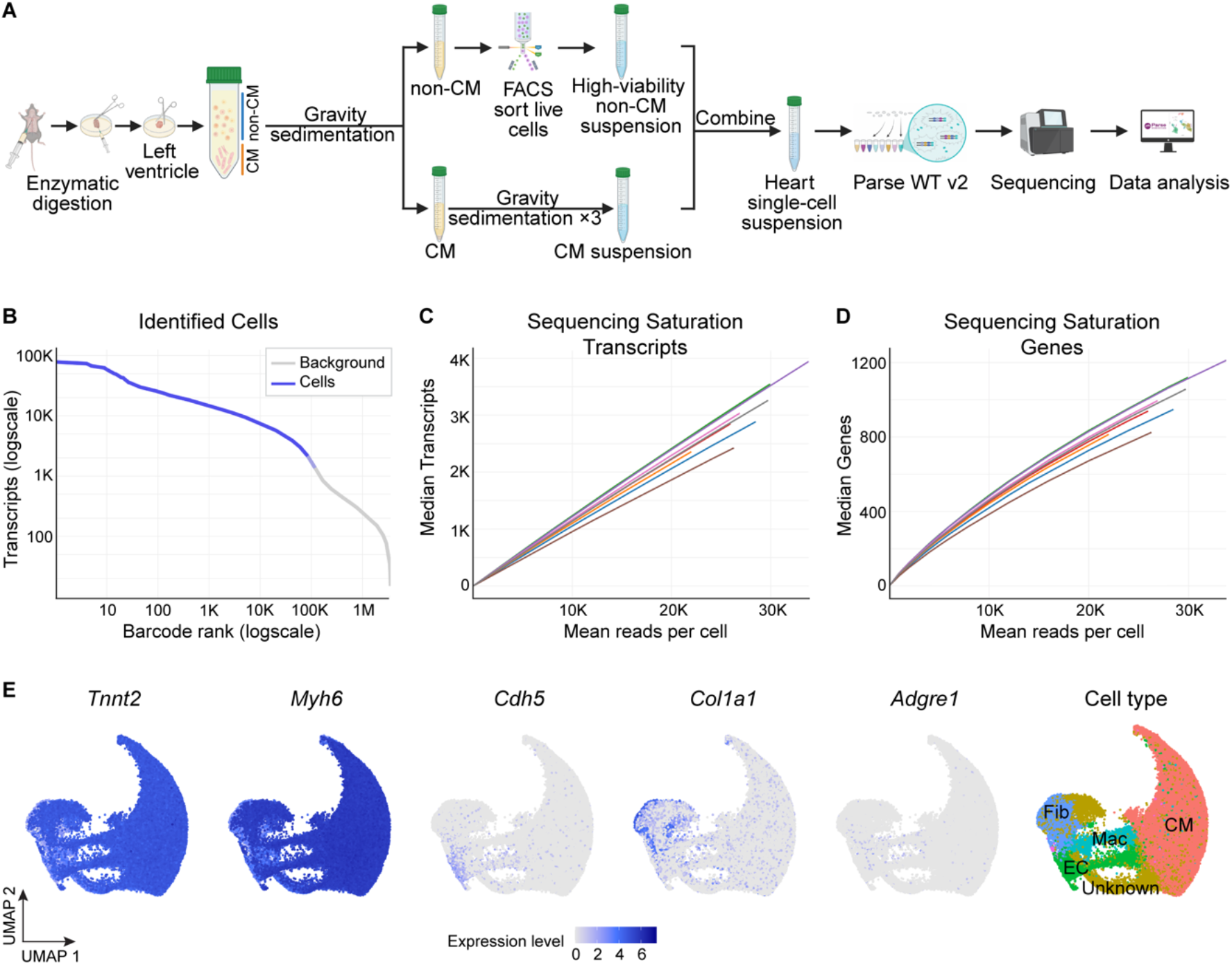
Mixing cardiomyocytes (CMs) with non-cardiomyocytes (non-CMs) skews single-cell libraries toward CMs. (A) Schematic for scRNA-seq of heart cells processed as a mixed CM + non-CM preparation. (B) Barcode-rank plot for all samples. (C–D) Sequencing-saturation curves for transcripts (C) and genes (D) for all samples; each curve represents one sublibrary. (E) Feature plots of canonical markers in the mixed libraries—cardiomyocytes (*Tnnt2, Myh6*), endothelial cells (*Cdh5*), fibroblasts (*Col1a1*), and macrophages (*Adgre1*), together with cell-type annotations. Fib, fibroblast; EC, endothelial cell; Mac, macrophage.

**Table 1.** Technical metrics across generated whole-cell and online nuclear libraries. Reads mapped to transcriptome: Fraction of reads with valid barcodes that map to an annotated gene (GTF file); Reads mapped to exonic regions: Fraction of reads with valid barcodes that map to an exon; Fraction reads in cell: Number of reads in cells divided by the total number of mapped reads; Slope of barcode-rank curve: Values that approach 5 and above have a significant “drop” at the cutoff point, while values that are closer to 1 do not show a clear transition between cell and background barcodes.

| Dataset | Mixed<br>lower depth | Mixed<br>higher depth | Separated<br>CMs | Separated<br>non-CMs | snRNA-seq |
| --- | --- | --- | --- | --- | --- |
| Estimated Number of Cells | 122,930 | 15,025 | 2,375 | 20,786 | 56,663 |
| Mean reads/cell | 27,727 | 83,843 | 260,799 | 38,287 | 25,211 |
| Median transcripts/cell | 2,941 | 7,776 | 64,657 | 9,092 | 1,840 |
| Median genes/cell | 965 | 1,968 | 4,881 | 2,869 | 1,202 |
| Sequencing saturation | 5.69% | 10.09% | 33.55% | 31.13% | 66.48% |
| Valid barcode | 57.47% | 52.75% | 89.17% | 96.56% | 96.25% |
| Reads mapped to transcriptome | 66.27% | 65.31% | 61.45% | 69.20% | 51.55% |
| Reads mapped to exonic regions | 77.41% | 77.41% | 67.40% | 44.10% | 20.90% |
| Fraction reads in cell | 38.65% | 38.31% | 86.15% | 84.07% | 64.85% |
| Slope of barcode-rank curve | 3.15 | 3.44 | 21.2 | 10.94 | NA |

Non-CMs were loaded in excess relative to CMs (∼2-fold) in the mixed-library experiment; however, downstream cell-type annotation revealed a strong bias toward CMs relative to the starting input ratio, with CMs outnumbering non-CMs by ∼4-fold (Figure 1E). Moreover, cells annotated as non-CMs showed unexpected expression of CM marker genes (e.g., *Myh6, Tnnt2*), which could result from CM ambient RNA contamination, and/or CM–non-CM multiplets (Figure 1E).

To test whether greater read depth could rescue sensitivity and recover more non-CMs, one sub-library was selected for re-sequencing at three-fold higher depth. Deeper sequencing modestly increased saturation and genes detected per cell but did not correct the absence of barcode-rank cliff (Figures S1A-C; Table 1), nor resolve the CM-skewed composition or CM-marker signal in the putative non-CM population (Figure S1D). Together, these results indicate that increasing sequencing depth alone is insufficient to salvage mixed CM+non-CM whole-cell libraries.

In summary, mixed libraries were unable to achieve high-quality output, characterised by lack of barcode-rank cliff configuration and under-representation of non-CMs despite non-CM-enriched input.

### CMs contain substantially more RNA and mitochondrial content than non-CMs

We hypothesised that the large size of CMs relative to non-CMs could influence the CM-dominant behavior of mixed libraries. Confocal imaging of isolated cells stained with Pyronin Y (RNA), MitoViewer (mitochondria), and Hoechst (nuclei) confirmed that adult CMs are substantially larger than non-CMs, with markedly higher mitochondrial and RNA content per cell (Figures 2A–2E). Independent bulk RNA quantification confirmed higher RNA yield per CM than per non-CM (Figure 2F). These results support a model wherein (1) high CM RNA content disproportionately contributes to library complexity and read allocation when CMs and non-CMs are co-processed, and (2) CMs are expected to require increased sequencing depth to achieve desirable saturation.

**Figure 2.**
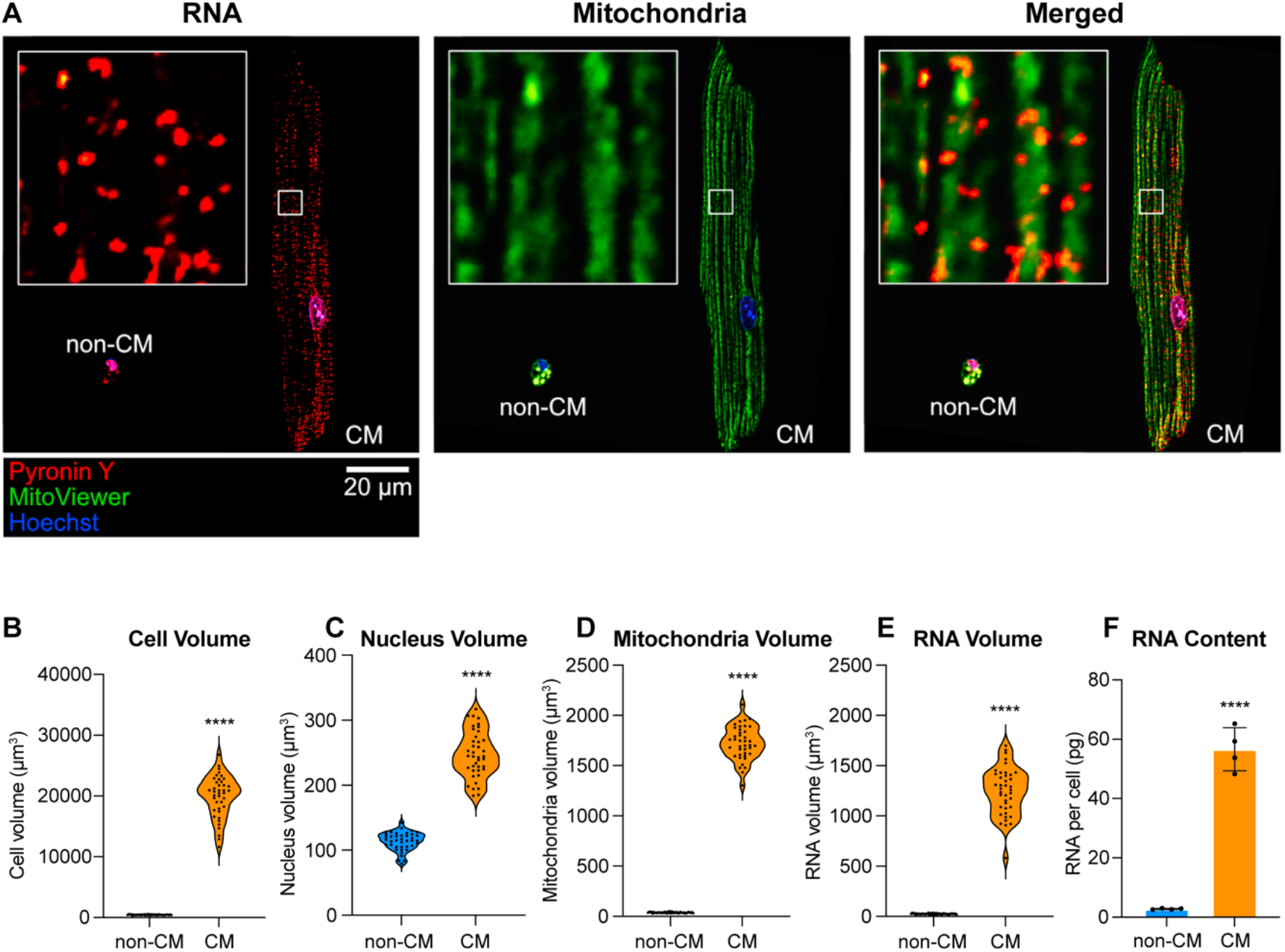
CMs contain substantially more RNA and mitochondria than non-CMs. (A) Representative confocal microscopy images of a non-CM and a CM. Cells were stained with Pyronin Y (red) for RNA, MitoViewer (green) for mitochondria, and Hoechst (blue) for nuclei. (B–E) Volumetric quantification of subcellular structures. (B) Cell volume, (C) Nucleus volume, (D) Mitochondrial volume, and (E) Total RNA volume were measured from confocal Z-stacks using Imaris surface rendering. CMs show significantly larger volumes across all parameters compared to non-CMs. Data points represent individual cells (n = 40). Statistical significance was determined by an unpaired, two-tailed Welch’s t-test. ****P < 0.0001. (F) Bulk RNA yield per cell: total RNA isolated from 10^5^ cells per sample using TRIzol extraction and quantified by NanoDrop. The RNA content per cell (pg) was calculated by dividing total RNA by input cell number. Data points represent independent mouse replicates (n = 4). Statistical significance was determined by an unpaired, two-tailed Welch’s t-test. ****P < 0.0001.

### Separating CMs from non-CMs with independent sequencing depth adjustment yields high-quality whole-cell CM scRNA-seq

Based on the above, we repeated our experiment, this time separating CMs and non-CMs before fixation/barcoding and processing them as independent libraries (Figure 3A). Focusing first on CM-only libraries, we observed the rescue of a distinctive barcode-rank cliff (Figure 3B), and sequencing saturation increased substantially relative to mixed libraries (Figures 3C–3D; Table 1). CM quality control (QC) metrics (transcripts per cell, genes detected per cell, mitochondrial RNA fraction) were consistent across samples (Figure 3E). For downstream visualisation and clustering, counts were normalised by per-cell library size and log-transformed. Canonical CM markers (*Tnnt2, Myh6, Ryr2*) were robustly detected, non-CM markers were largely absent, and rare non-CM impurities could be straightforwardly identified as an isolated cluster (Figure 3F). Once again, non-CM impurities in the CM fraction were found to express comparable levels of CM markers, alongside endothelial (EC) markers (*Cdh5, Pecam1*), and are likely to represent cell multiplets. Thus, CM isolation followed by CM-specific library construction and increased sequencing depth supports high-quality whole-cell CM scRNA-seq.

**Figure 3.**
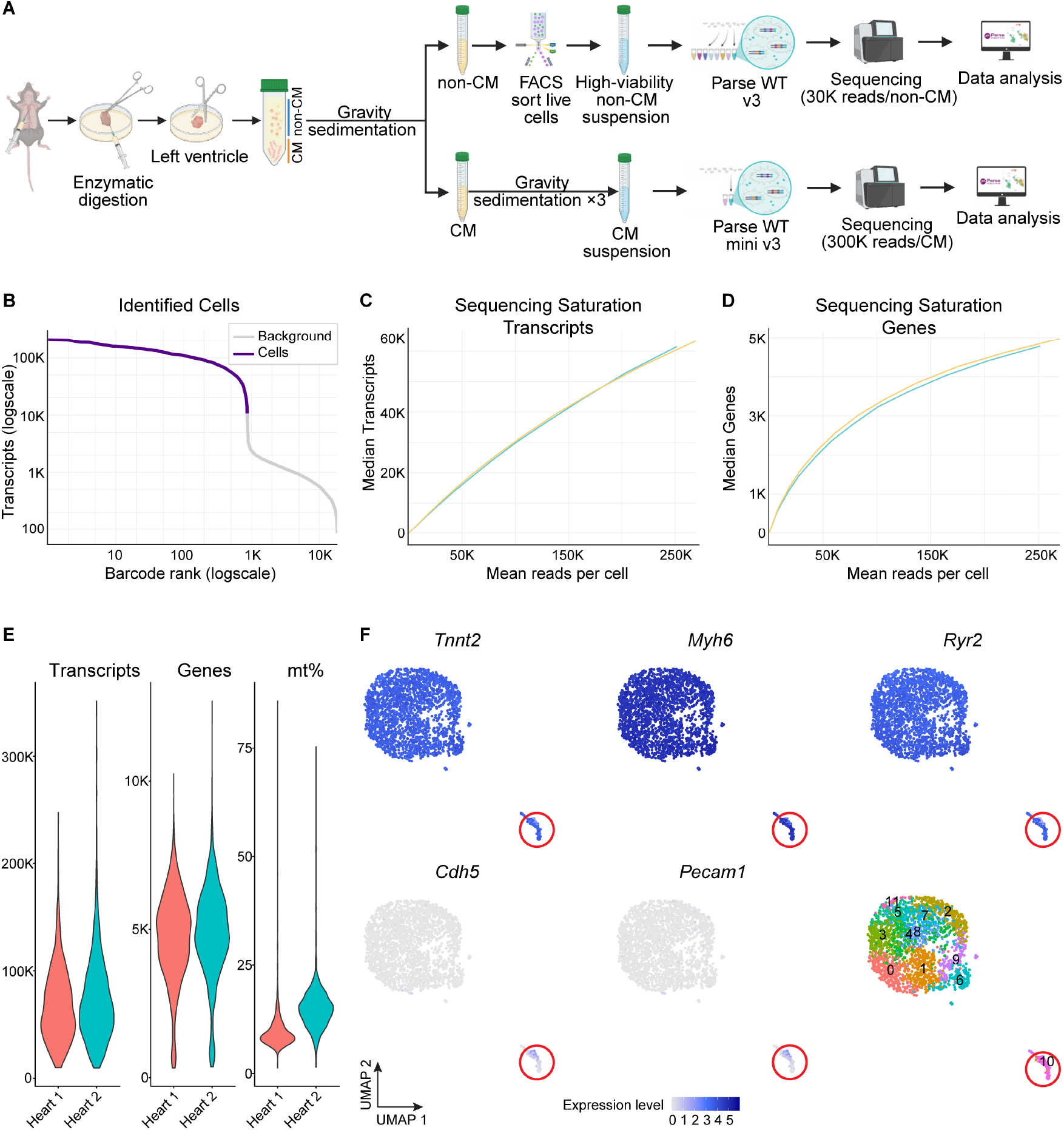
Optimisation enables high-quality whole-cell cardiomyocyte scRNA-seq. (A) Workflow schematic: CMs and non-CMs are processed in separate libraries; CM libraries are sequenced at increased depth. (B) Barcode-rank plot for a representative CM-only sample. (C–D) Sequencing-saturation curves for transcripts (C) and genes (D) in CM-only libraries; each curve represents one sublibrary. (E) Quality control (QC) summaries for CMs: transcripts per cell (left), genes per cell (middle), and mitochondrial RNA fraction (mt%; right). (F) Feature plots of canonical markers in the CM library—CMs (*Tnnt2, Myh6, Ryr2*) and endothelial cells (*Cdh5, Pecam1*), together with Seurat cluster annotations. Red circles highlight cluster 10, a small contaminating endothelial cell population.

### Concomitant high-quality non-CM scRNA-seq with broad cell-type recovery

Non-CM-only libraries also exhibited a clear barcode-rank cliff (Figure 4A), increased sequencing saturation (Figures 4B and 4C; Table 1), and consistent per-cell QC metrics across samples (Figure 4D). Unsupervised clustering resolved major cardiac non-CM populations (Figure 4E), supported by canonical marker expression (Figure 4F) and top differentially expressed markers for each annotated cell type (Figure 4G). Importantly, CM markers were largely absent, suggesting low ambient RNA capture, and rare CM impurities, possibly comprising small CMs, could be straightforwardly identified as a well-separated cluster (Figure 4F).

**Figure 4.**
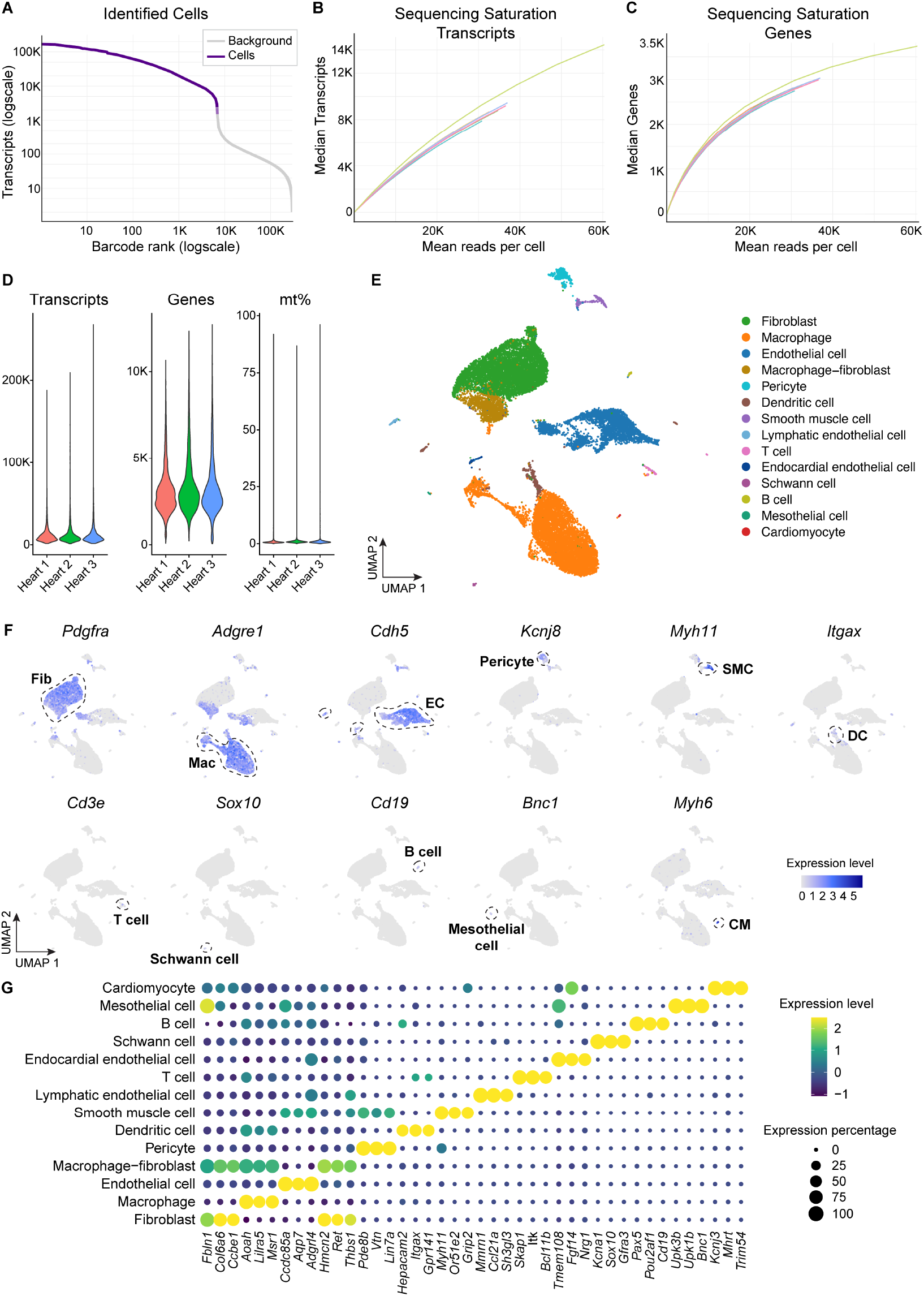
Optimisation yields high-quality scRNA-seq of heart non-cardiomyocytes. (A) Barcode-rank plot for a representative non-CM-only sample. (B–C) Sequencing-saturation curves for transcripts (B) and genes (C) in non-CM libraries; each curve represents one sublibrary. (D) QC summaries for non-CMs: transcripts per cell (left), genes per cell (middle), and mitochondrial RNA fraction (mt%; right; generally lower than CMs in Fig. 3). (E) Integrated UMAP of non-CMs colored by annotated cell type. (F) Feature plots of canonical markers for major non-CM lineages — fibroblasts (*Pdgfra*), macrophages (*Adgre1*), endothelial cells (*Cdh5*), pericytes (*Kcnj8*), smooth muscle cells (*Myh11*), dendritic cells (*Itgax*), T cells (*Cd3e*), Schwann cells (*Sox10*), B cells (*Cd19*), mesothelial cells (*Bnc1*), together with the CM marker *Myh6*, which identifies a small contaminating CM population. Dashed circles indicate the corresponding populations. Fib, fibroblast; EC, endothelial cell; Mac, macrophage; SMC, smooth muscle cell; DC, dendritic cell. (G) Dot plot of top marker genes per cell type (expressed in ≥50% of cells; FDR < 0.05; ranked by average log2 fold change).

Gene biotype composition also differed between the separated whole-cell CM and non-CM libraries. The fractions of detected genes were broadly similar between CM and non-CM libraries, and were dominated by protein-coding genes (Figure S2A). However, the distribution of total expression differed: non-CM libraries were dominated by protein-coding transcripts, whereas CM libraries contained a much larger fraction of rRNA and mitochondrial-encoded transcripts (Figure S2B). Within the rRNA pool, nuclear rRNA-related genes accounted for the majority of reads in both CM and non-CM libraries, while mitochondrial rRNAs (*mt-Rnr1* and *mt-Rnr2*) contributed a larger share in CMs than in non-CMs (Figures S2C and S2D). These data are consistent with the elevated rRNA and mitochondrial content in CMs.

### Comparison with snRNA-seq highlights the complementary strengths of whole-cell scRNA-seq

Because snRNA-seq is widely used for adult CM transcriptomics, an important question was what additional information whole-cell scRNA-seq could provide. We therefore compared our whole-cell CM and non-CM datasets to a recently published mouse left-ventricle snRNA-seq dataset(13).

Technical metrics differ between whole-cell and nuclear libraries. Across datasets, whole-cell libraries were generated at higher mean reads per cell, whereas snRNA-seq reached higher sequencing saturation at lower sequencing depth (Table 1). This pattern is consistent with higher library complexity in whole-cell preparations, in which reads are distributed across a larger pool that includes cytoplasmic and mitochondrial transcripts and therefore require deeper sequencing to approach saturation. Accordingly, additional sequencing would be expected to continue to recover more transcripts/genes in scRNA-seq. In addition, whole-cell data showed a higher fraction of reads mapping to the transcriptome and to exonic regions (Table 1), consistent with capture of mature cytoplasmic mRNAs, while nuclear RNA libraries are expected to contain a larger proportion of unspliced/intronic reads. Finally, the fraction of reads assigned to cells was higher in our separated whole-cell libraries (Table 1), suggesting reduced background relative to the snRNA-seq dataset.

To define the biological consequences of the technical differences, we next compared cardiomyocyte expression profiles between whole-cell scRNA-seq and snRNA-seq, using a pseudobulk DESeq2 framework followed by gene ontology (GO) Biological Process enrichment (Figure 5). Genes up-regulated in whole-cell cardiomyocyte scRNA-seq were enriched for processes related to cellular respiration, cytoplasmic translation, and heart contraction (Figure 5A), whereas genes up-regulated in cardiomyocyte snRNA-seq were enriched primarily for processes related to regulation of gene expression (Figure 5B). Representative differentially expressed genes illustrating these method-dependent biases are shown in Figure 5C. Together, these results show that the value of whole-cell CM profiling lies not only in recovering more RNA overall, but also in preferential recovery of metabolic, cytoplasmic translation, and contractile gene programs that are under-represented in nuclear-only profiling.

**Figure 5.**
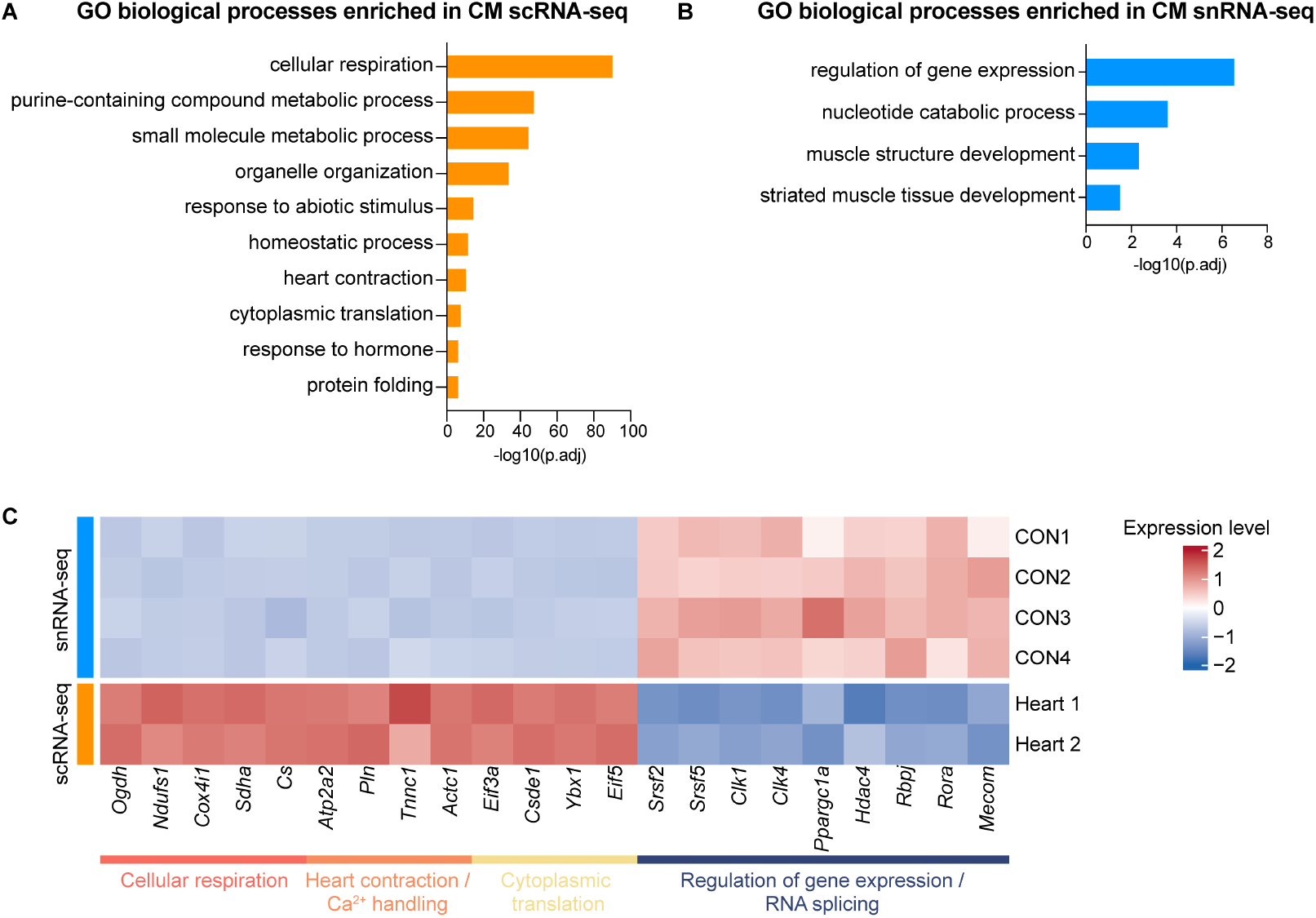
Comparison of cardiomyocyte whole-cell scRNA-seq and snRNA-seq. (A) Pathway enrichment of genes up-regulated in whole-cell CM scRNA-seq. Differential expression between whole-cell CM scRNA-seq and snRNA-seq was performed using pseudobulk profiles and DESeq2. Genes up-regulated in whole-cell CM scRNA-seq were defined as log_2_ fold change > 1, adjusted p-value < 0.001, and baseMean > 1500. GO Biological Process enrichment was performed using g:Profiler, and the top 10 enriched driver terms are shown. (B) Pathway enrichment of genes up-regulated in CM snRNA-seq. Genes up-regulated in CM snRNA-seq were defined as log_2_ fold change < −1, adjusted p-value < 0.001, and baseMean > 1500, using the same pseudobulk DESeq2 framework as in (A). GO Biological Process enrichment was performed using g:Profiler, and the enriched driver terms are shown. (C) Heatmap of representative differentially expressed genes between whole-cell CM scRNA-seq and snRNA-seq identified by pseudobulk DESeq2 analysis. Genes are grouped by functional categories, highlighting enrichment of cellular respiration, heart contraction/Ca^2+^ handling, and cytoplasmic translation genes in whole-cell CM scRNA-seq, and enrichment of regulation of gene expression/RNA splicing genes in snRNA-seq.

Non-cardiomyocyte comparisons show heterogeneous effects. We performed analogous analyses in major non-cardiomyocyte lineages (fibroblasts, endothelial cells, and macrophages) using pseudobulk-based differential expression followed by gene-set enrichment analysis (GSEA; Figure S3). Across these cell types, compared to snRNA-seq, whole-cell scRNA-seq tended to show enrichment of cytoplasmic processes, but features displayed cell-type specificity. Fibroblast scRNA-seq enrichment included terms associated with cellular export and extracellular matrix proteins, endothelial cells with membrane-associated functions such as phagocytosis, cell adhesion, and immune interactions, and macrophages with immune function but also, interestingly, chromatin organisation and post-transcriptional gene regulation. In contrast, in all three non-CM lineages, snRNA-seq was associated with enrichment of CM-associated programs, which is consistent with previously reported technical limitations in snRNA-seq methodology resulting in increased residual ambient background CM RNA, rather than bona fide non-CM biology(2). These observations suggest that method-driven differences in non-cardiomyocyte comparisons may be confounded by background signal and should be interpreted with caution.

## DISCUSSION

Adult CMs pose a distinctive challenge for whole-cell scRNA-seq because they are unusually large and fragile. In this study, we developed a practical high-quality and scalable workflow for whole-cell scRNA-seq of adult hearts. We identified per-cell RNA content as a core consideration that should be accounted for to prevent failure in mixed-library designs, due to complexity and domination by large, high RNA-content cell types, which could not be rescued by increased sequencing depth. By separating CMs and non-CMs prior to library construction and allocating sequencing depth according to RNA content, we obtained high-quality whole-cell CM and non-CM datasets from the same heart preparation. More broadly, this study establishes a general workflow and guiding principles for samples containing large cells or cell populations with highly unequal RNA content.

The workflow herewith provides a high-quality and scalable approach for whole-cell scRNA-seq of adult CMs, expanding the currently limited set of available methods. It has been at times presumed that very high (80%) mitochondrial transcript fractions observed when subjecting CMs to simple FACS scRNA-seq approaches are attributable to the biological reality of high mitochondrial content in these metabolically active cells. However, we found that although increased relative to non-CMs (Figure S2B), mitochondrial transcript fractions remained largely below 25% (Figure 3E). This is more in line with lower throughput, improved large particle FACS, which measured 40%, similar to bulk RNA-seq(7), 30% measured by Mercer et al. in total heart RNA(14), and high-quality manual cell-picking approaches(11, 12). Interestingly, our re-analysis of a published ICELL8 dataset(10) showed a high mitochondrial transcript fraction in the CM population (Figure S4), suggesting a potential challenge in CM data quality using this approach. Furthermore, such nanowell-based platforms require stringent image-based quality control and typically recover only hundreds of intact CMs per chip(10, 15). It is important to highlight that mitochondrial transcripts, rather than simply a QC metric, are also biologically informative for questions concerning cell metabolism, mitochondria, and energetic stress, and accurate quantification is therefore in itself meaningful.

The technical metrics in Table 1 are informative for interpreting data quality. Sequencing saturation should be interpreted in the context of library complexity: whole-cell CM libraries saturate more slowly than non-CM libraries because each CM contains a larger and more complex transcript pool. Similarly, whole-cell libraries in general saturate more slowly than snRNA-seq libraries because they include cytoplasmic and mitochondrial transcripts in addition to nuclear RNA. Accordingly, whole-cell scRNA-seq typically requires deeper sequencing to approach saturation. Therefore, even with relatively higher reads per cell in scRNA-seq, sequencing saturation may remain low, but additional sequencing has greater potential to achieve more transcripts/genes per cell. Higher fractions of reads mapped to the transcriptome and exonic regions are expected in whole-cell libraries, which capture more mature cytoplasmic mRNAs, whereas snRNA-seq yields a larger fraction of intronic and pre-mRNA reads(3). The marked increase in reads-in-cells fraction and barcode-rank slope after separating CMs and non-CMs are evidence of reduced background and improved cell calling. Differences in valid barcode fraction are likely influenced by both workflow design and manufacturer product chemistry version, which at time of writing is continually being updated, and should therefore be interpreted cautiously.

The comparison between whole-cell CM scRNA-seq and CM snRNA-seq further highlights that these methods are complementary rather than interchangeable. In our analysis, whole-cell CM scRNA-seq profiles were enriched for cellular respiration, cytoplasmic translation, and contractile programs, whereas snRNA-seq preferentially emphasised gene regulation signatures (Figure 5). This is consistent with the broader understanding that whole-cell scRNA-seq captures cytoplasmic transcripts more effectively, while snRNA-seq provides a more nucleus-centered view of transcriptional regulation. Practical method choice should therefore be guided by the biological question. For studies focused on CM metabolism, mitochondria, cytoplasmic translation, or contractile biology, whole-cell CM scRNA-seq is likely to be the more informative method. By contrast, snRNA-seq remains powerful for frozen/stored or hard to dissociate tissues, and questions centered on nuclear transcriptional regulation.

In contrast to mitochondrial transcript detection, the elevated rRNA fraction in CM library (Figure S2B) is a practical limitation of whole-cell CM scRNA-seq. rRNA contributes relatively little desired information for most downstream analyses, while consuming substantial sequencing capacity. Future wet-lab optimisation of whole-cell CM scRNA-seq workflows might further include strategies to reduce rRNA capture, or alternatively, plan sequencing depth with this cost in mind. Loi et al. have utilised rRNA depletion to enrich the remaining transcriptome in single-cell total RNA-seq(16), supporting the feasibility and utility of this direction.

Our non-CM dataset highlights a limitation of enzymatic dissociation-based single-cell workflows: recovered cell-type proportions do not necessarily reflect *in vivo* abundance. In the adult mouse heart, endothelial cells are generally understood to constitute the majority of non-CMs, whereas fibroblasts and myeloids are a smaller fraction(17). In our workflow, endothelial cells were under-represented, while macrophages and fibroblasts were relatively over-represented. This likely reflects a combination of dissociation bias, differential cell fragility, and viability selection. Such biases are not unique to the heart. Enzymatic dissociation is known to induce stress-response(2, 3, 18), and method-dependent differences in cell-type recovery and biological interpretation have also been observed in other organs. For example, in kidney, scRNA-seq better captured immune cells, whereas snRNA-seq provided superior insight into tubular and epithelial cells(3, 19); in liver, scRNA-seq detected more immune cells, whereas snRNA-seq preferentially recovered hepatocytes and endothelial cells(18, 20). Therefore, non-CM cell-type composition in our dataset should be interpreted as workflow-dependent recovery rather than a direct estimate of physiological abundance.

Low-level impurities in the separated libraries were expected. In the CM library, minor endothelial cell contamination is consistent with close CM-EC association resulting in EC impurities in isolated CM fractions in previous reports(21). In our dataset, these EC impurities were present at low abundance, formed a readily distinguishable cluster, and could be removed straightforwardly before downstream analyses. In the non-CM library, the CM cluster may reflect a population of CMs small enough to filter into the non-CM sorting gate. These impurity populations were minor and readily distinguishable by canonical marker expression, and could also be removed straightforwardly before downstream analyses.

Methodologically, in summary, we recommend several practice principles in whole-cell scRNA-seq. First, when a sample contains cell populations with dramatically different RNA content, such as CMs and non-CMs, these populations should be processed as separate libraries rather than co-processed in a pooled library. Second, in such cases, sequencing depth should be allocated according to expected per-cell RNA content rather than evenly across populations. In our experience, adult whole-cell CMs required substantially deeper sequencing, approximately 300,000 reads per cell, whereas non-CMs were sequenced at approximately 30,000 reads per cell. Third, method choice should follow the biological question: whole-cell CM scRNA-seq is advantageous for metabolism, mitochondria, cytoplasmic translation, and contraction centered studies, whereas snRNA-seq remains powerful for frozen/stored tissues and nucleus centered gene regulation studies. More broadly, our whole-cell scRNA-seq workflows and principles can be extended beyond the adult heart. Candidate contexts include skeletal muscle cells, hepatocytes, megakaryocytes, large neurons, and other potentially large or RNA-rich cell types, although these applications may require context-specific validation.

### Limitations of the study

This study has several limitations. First, our comparison between whole-cell scRNA-seq and snRNA-seq is a cross-platform comparison, and therefore the observed differences are not attributed solely to cell-versus-nucleus biology, as they may also reflect differences in chemistry, alignment, counting strategies, and sequencing depth. In addition, differential expression between whole-cell and nuclear datasets should be interpreted as reflecting the relative transcript proportion in the two methods, rather than absolute transcript quantification, because nuclei capture only a subset of total cellular RNA and whole-cell libraries are therefore expected to contain a larger total transcript pool than nuclear libraries. Second, as discussed above, non-CM recovery in the whole-cell workflow does not fully preserve physiological abundance. Third, the high rRNA fraction in whole-cell CM library reduces sequencing efficiency and remains a target for future optimisation. Future work should include matched cell-versus-nucleus profiling from the same tissue, with matched-depth and matched-annotation, and experimental strategies to reduce rRNA capture. Taken together, our findings do not aim to position whole-cell CM scRNA-seq as superior or a replacement for snRNA-seq; rather, they define when whole-cell CM profiling might offer a beneficial choice, and technical suggestions for successful conduct.

## RESOURCE AVAILABILITY

### Lead contact

Requests for further information and resources should be directed to and will be fulfilled by the lead contact, Matthew Andrew Ackers-Johnson.

### Materials availability

This study did not generate new unique reagents.

### Data and code availability

Accession numbers and code deposition will be finalised before journal publication.

## ACKNOWLEDGMENTS

The authors thank Xiaoning Wang and Delia Pang from the NUS Medicine Flow Cytometry Laboratory for assistance and advice on cell sorting and analysis. The authors acknowledge the following funding sources: Singapore Ministry of Education Grant MOET32021-0003 (M.A.-J. and R.S.Y.F.), and the Singapore Ministry of Health National Medical Research Council (NMRC) Clinician Scientist Senior Investigator Award MOH-001685 (R.S.Y.F.)

## AUTHOR CONTRIBUTIONS

Conceptualisation, M.A.-J.; Methodology, M.A.-J., Y.H., R.G., J.S., N.R.; Investigation, Y.H., M.A.-J., S.M., T.D.A.L., J.S., S.N., B.Y.C.T.; Formal analysis, Y.H. and E.V.; Data curation, Y.H.; writing—original draft, Y.H. and M.A.-J.; writing—review & editing, H.C., B.L., H.Y.; Funding acquisition, M.A.-J and R.S.Y.F.; Supervision, M.A.-J and R.S.Y.F.

## TABLES AND TEXT BOXES

## SUPPLEMENTAL INFORMATION

Supplementary Figure titles and legends:

**Figure S1.**
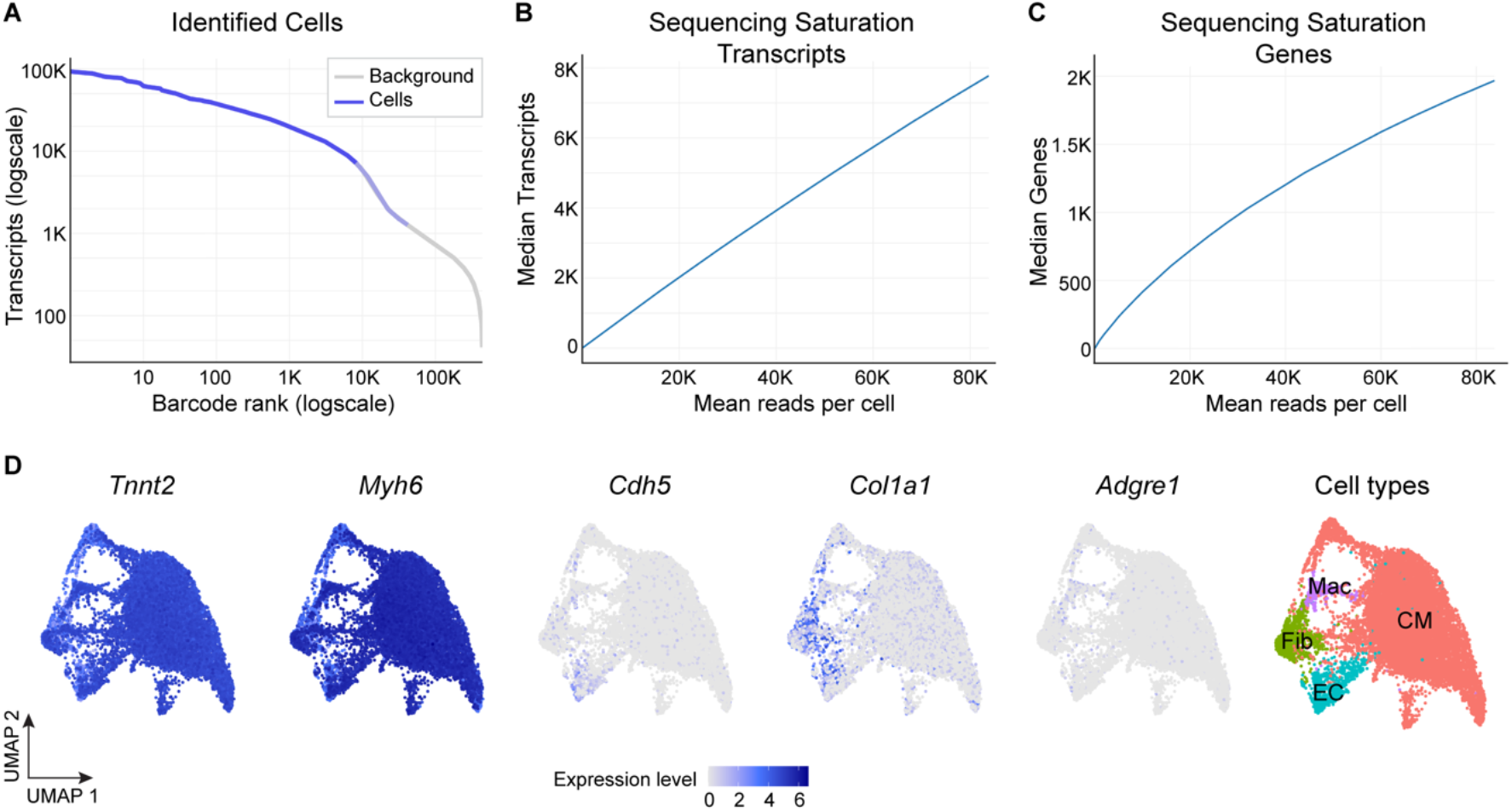
Increasing sequencing depth for mixed library is insufficient to salvage data. (A-C) Deeper resequencing of one sublibrary: barcode-rank (A) and sequencing-saturation (B, C). (D) Feature plots of the canonical markers after deeper resequencing of the sublibrary – cardiomyocytes (*Tnnt2, Myh6*), endothelial cells (*Cdh5*), fibroblasts (*Col1a1*), and macrophages (*Adgre1*), together with cell-type annotations. Additional depth increased sensitivity but did not correct the CM-skewed cell-type composition introduced by co-processing. Fib, fibroblast; EC, endothelial cell; Mac, macrophage.

**Figure S2.**
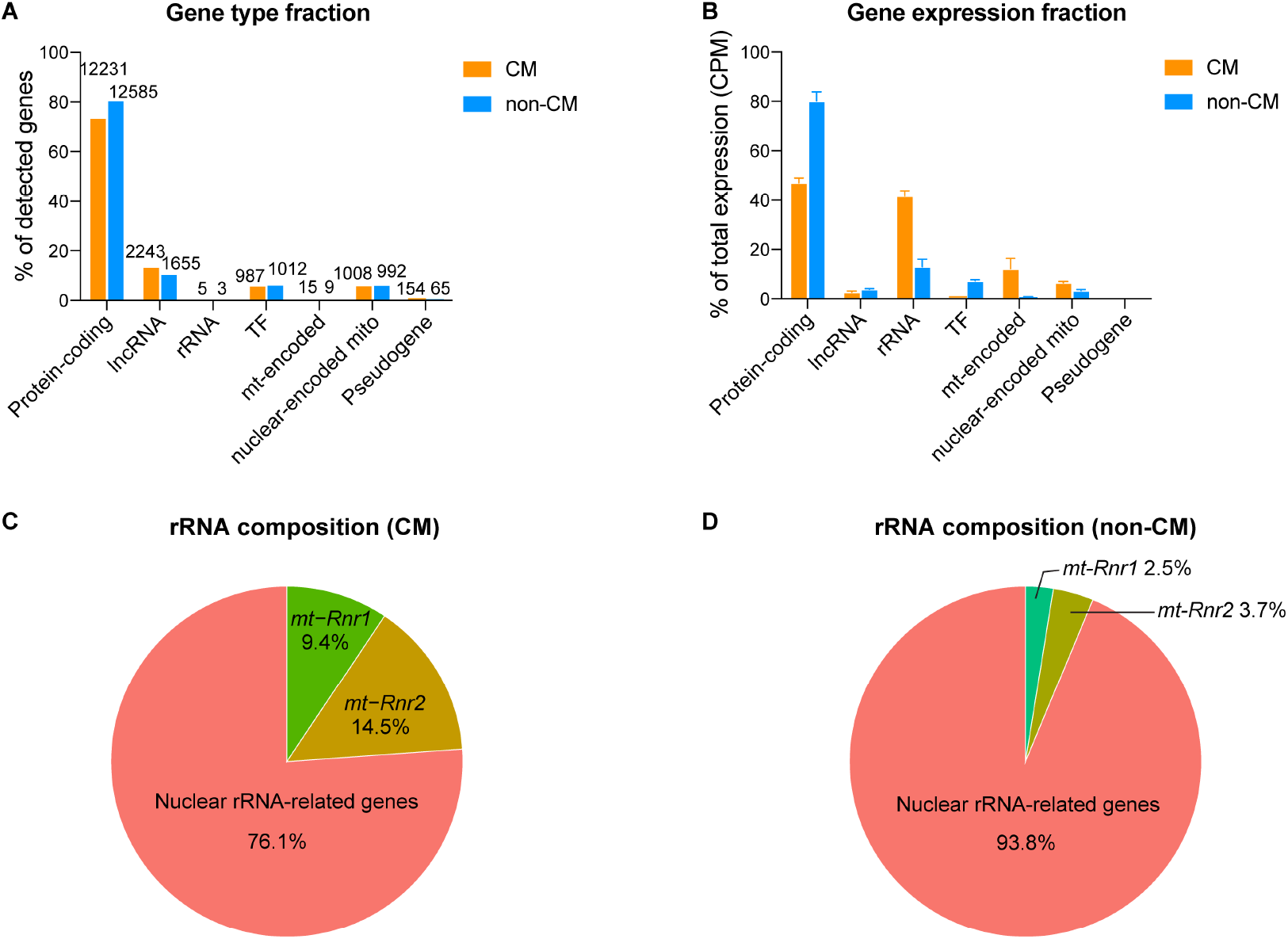
Gene-class composition in whole-cell scRNA-seq of cardiomyocytes and non-cardiomyocytes. (A) Gene-class detection in CMs and non-CMs. Bar plot showing the percentage of detected genes belonging to each gene class (protein-coding, long non-coding RNA (lncRNA), transcription factors (TFs), mitochondrial-encoded genes (mt-encoded), nuclear-encoded mitochondrial genes (nuclear-encoded mito), rRNA, and pseudogenes) in whole-cell CM scRNA-seq. A gene was considered detected if expressed in ≥ 1% of cells in each dataset. Because gene classes overlap (e.g., TFs and nuclear-encoded mitochondrial genes are largely protein-coding), gene classes are non-exclusive, and therefore percentages do not sum to 100%. Numbers above each bar indicate the number of detected genes in the corresponding gene class. (B) Gene-class expression composition in CMs and non-CMs. Bar plot showing the fraction of total expression attributable to each gene class, calculated from pseudobulk CM and non-CM profiles. For each sample, counts were aggregated across CM cells, normalised to counts per million (CPM), and summarised as: fraction = (sum of CPM for genes in a given class) / (sum of CPM for all genes in the count matrix). (C) rRNA expression composition within rRNA genes in CM scRNA-seq. Pie chart showing the relative contribution of individual rRNA genes to total rRNA expression in CMs scRNA-seq. For each sample, counts were aggregated across CMs to generate pseudobulk matrix, normalised to counts per million (CPM), and rRNA fractions were calculated within the rRNA gene set as: fraction of gene_i = CPM _i / (sum CPM of all rRNA genes). Fractions were computed per sample and then averaged across samples; pie slices indicate the mean percentage. (D) rRNA expression composition within rRNA genes in non-CMs scRNA-seq. Pie chart showing the relative contribution of individual rRNA genes to total rRNA expression in non-CMs scRNA-seq, calculated analogously to (C) using pseudobulk CPM per sample and summarizing expression within the rRNA gene set, followed by averaging across samples. Pie slices indicate the mean percentage.

**Figure S3.**
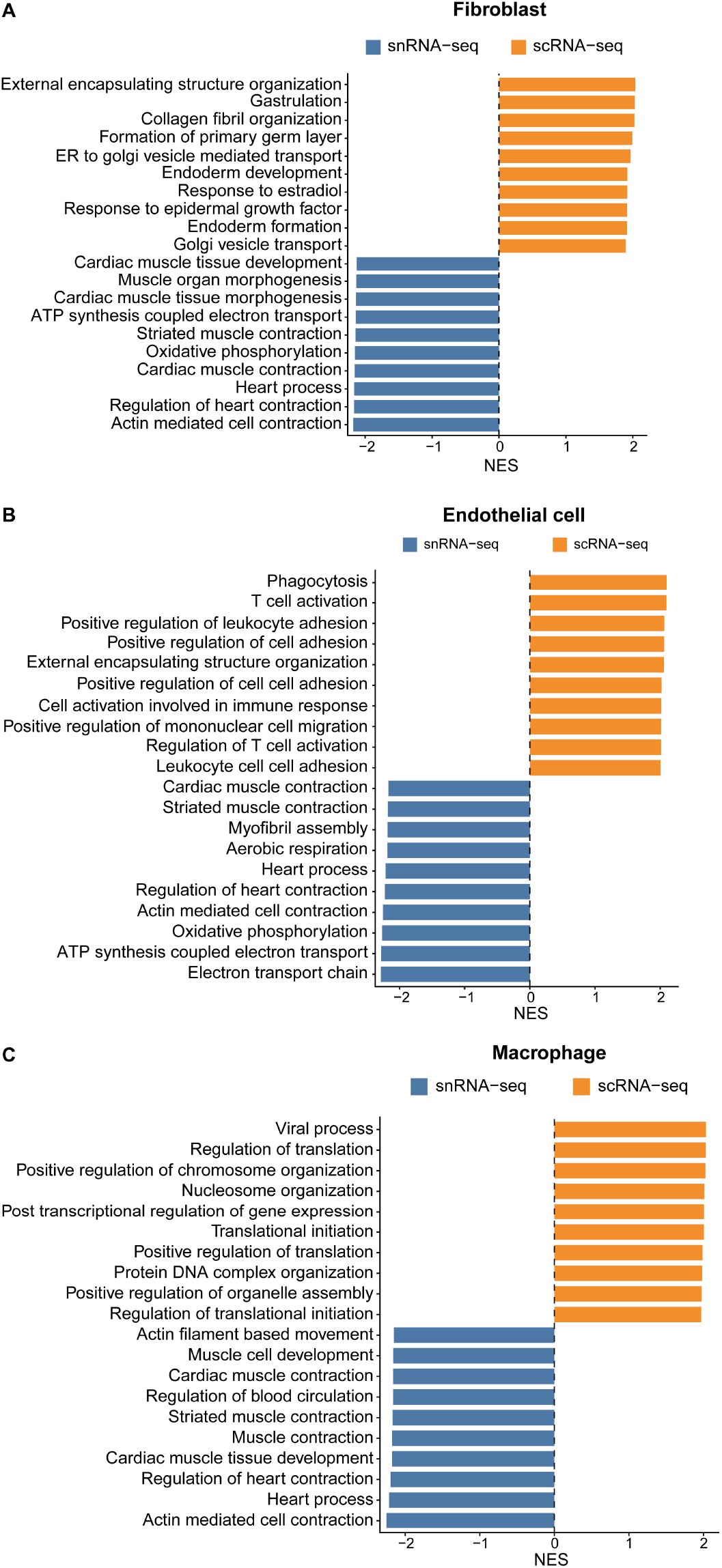
Comparison of non-cardiomyocyte whole-cell scRNA-seq and snRNA-seq. (A–C) Enrichment analysis highlights biological processes differentially enriched between whole-cell scRNA-seq and snRNA-seq across major non-CM cell types. Differential expression was performed within each cell type (A, fibroblast; B, endothelial cell; C, macrophage) using a pseudobulk strategy. For each dataset, raw counts were aggregated per sample across cells of the indicated cell type to generate pseudobulk matrices. Differential expression between datasets was calculated using DESeq2, comparing scRNA-seq versus snRNA-seq. Genes were ranked by −log_10_(p value) × sign(log_2_ fold change), and GSEA was performed using clusterProfiler with GO Biological Process gene sets (msigdbr, Mus musculus). Bar plots show the top 10 positively enriched pathways (NES > 0; enriched in scRNA-seq; orange) and top 10 negatively enriched pathways (NES < 0; enriched in snRNA-seq; blue) for each cell type. NES, normalised enrichment score.

**Figure S4.**
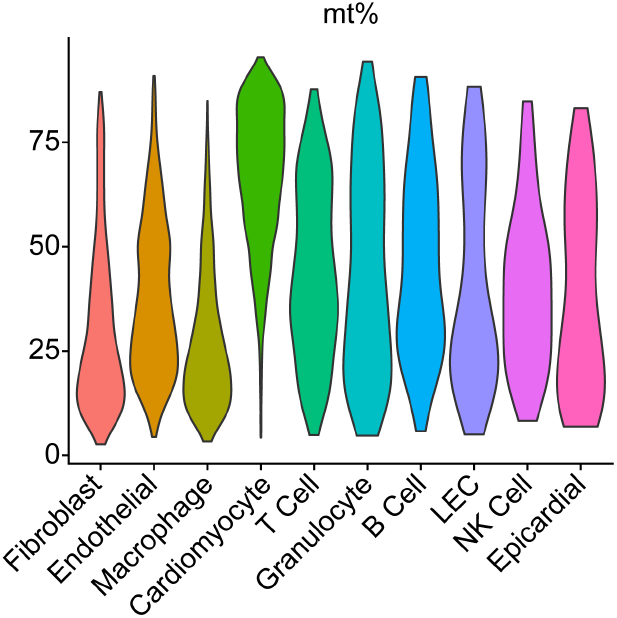
Re-analysis of a published ICELL8 dataset shows a high mitochondrial transcript fraction in cardiomyocytes. Violin plot shows the distribution of mitochondrial transcript fraction (mt%) across cell populations in the published ICELL8 dataset(10). CMs exhibited a higher mt%, highlighting a potential challenge for maintaining cardiomyocyte data quality with this approach. LEC, lymphatic endothelial cell; NK, natural killer cell.

**Figure S5.**
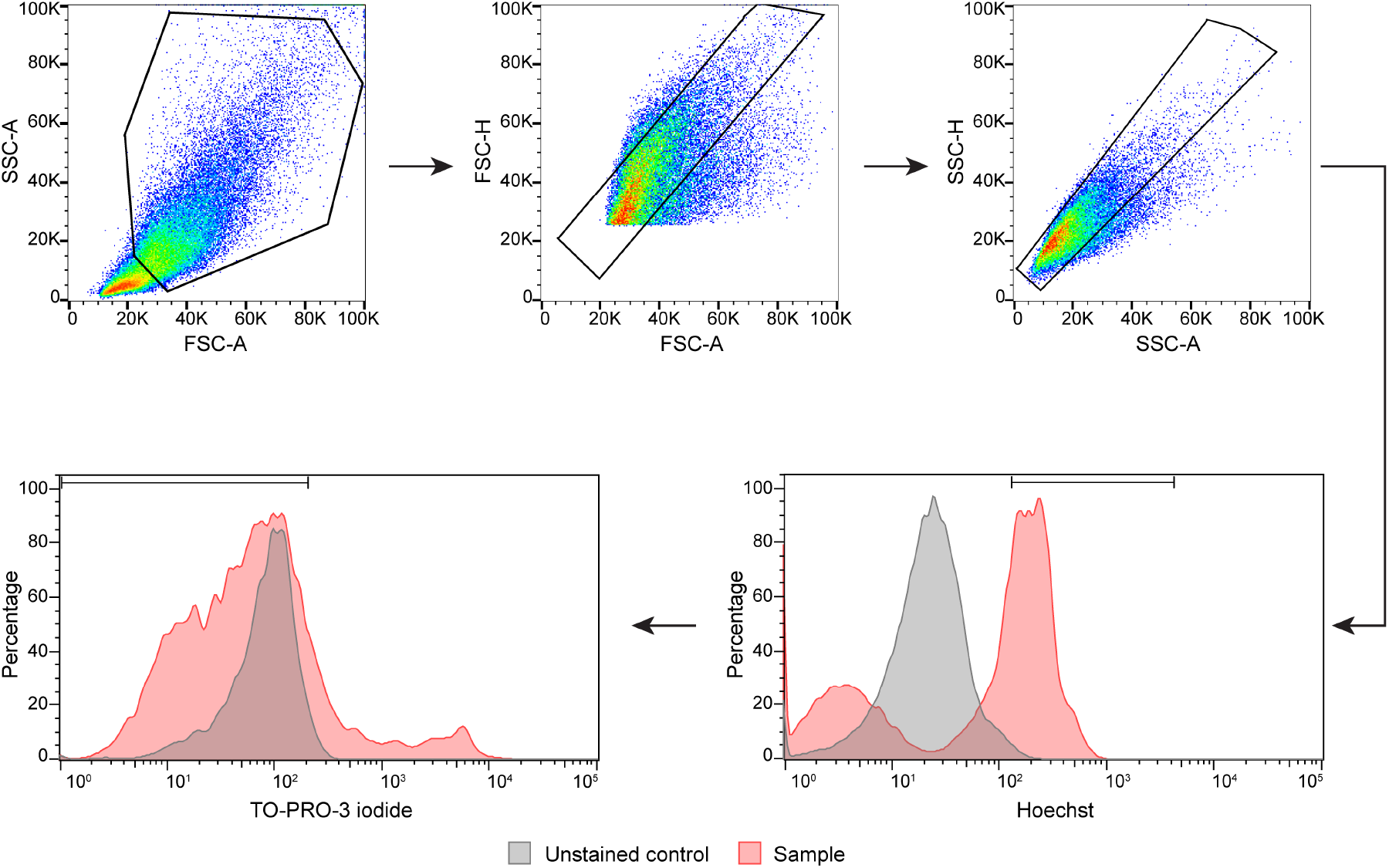
Gating strategy for sorting single live non-cardiomyocytes by FACS. Non-CMs were stained with Hoechst and TO-PRO-3 iodide. Single live non-CMs were defined as Hoechst-positive and TO-PRO-3 iodide-negative.

## STAR★METHODS

## KEY RESOURCES TABLE

| REAGENT OR RESOURCE | SOURCE | IDENTIFIER |
| --- | --- | --- |
| Antibodies |  |  |
| - | - | - |
| Bacterial and virus strains |  |  |
| - | - | - |
| Biological samples |  |  |
| Adult mouse heart tissues | C57BL/6 | For freshly isolated CMs and non-CMs; see Method Details |
| Chemicals, peptides, and recombinant proteins |  |  |
| TO-PRO-3 iodide | Thermo Fisher Scientific | T3605 |
| Hoechst 33342 | Thermo Fisher Scientific | H3570 |
| Pyronin Y | Sigma Aldrich | P9172 |
| MitoViewer (MitoView Green) | Biotium | 70054 |
| TRIzol | Thermo Fisher Scientific | 15596018 |
| Collagenase II | Worthington | LS004177 |
| Collagenase IV | Worthington | LS004189 |
| Dispase | Thermo Scientific | 17105041 |
| ActD | Sigma Aldrich | A9415 |
| Proteinase XIV | Sigma Aldrich | P5147 |
| DNase I | Thermo Scientific | 18047019 |
| Bovine Albumin Fraction V (7.5% solution) | Thermo Scientific | 15260037 |
| Critical commercial assays |  |  |
| Evercode™ Cell Fixation v2 | Parse Biosciences |  |
| Evercode™ WT v2 | Parse Biosciences |  |
| Evercode™ Cell Fixation v3 | Parse Biosciences |  |
| Evercode™ WT Mini v3 | Parse Biosciences |  |
| Evercode™ WT v3 | Parse Biosciences |  |
| Deposited data |  |  |
| Mouse left-ventricle snRNA-seq dataset | Li et al., 2025 | GEO: GSE263648 |
| Published mouse heart ICELL8 scRNA-seq dataset | Ren et al., 2020 | GEO: GSE120064 |
| Experimental models: Cell lines |  |  |
| - | - | - |
| Experimental models: Organisms/strains |  |  |
| Mus musculus, C57BL/6J |  | males, 3 months old |
| Oligonucleotides |  |  |
| - | - | - |
| Recombinant DNA |  |  |
| - | - | - |
| Software and algorithms |  |  |
| split-pipe | Parse Biosciences | v1.1.2 (pilot); v1.6.0 (optimized workflow) |
| R | The R Project | 4.5.1 |
| Seurat | Satija Lab | 5.4.0 |
| Other |  |  |
| NovaSeq X Plus | Illumina |  |
| NanoDrop spectrophotometer | Thermo Fisher Scientific |  |

## EXPERIMENTAL MODEL AND STUDY PARTICIPANT DETAILS

### Animals

Adult male C57BL/6J mice aged 3 months were used for all experiments reported in this study. Mice were maintained under standard housing conditions with *ad libitum* access to food and water and a 12 h light/12 h dark cycle. All mouse work was performed in accordance with Singapore National Advisory Committee for Laboratory Animal Research guidelines and approved by the NUS Institutional Animal Care and Use Committee.

## METHOD DETAILS

### Adult heart dissociation and isolation of cardiomyocytes and non-cardiomyocytes

Adult ventricular cells were isolated as previously described by Ackers-Johnson et al.(22), with modifications suitable for subsequent cell fixation and split-pool barcoding. Hearts were digested to generate a whole-cell suspension that was pre-stained with Hoechst in digestion buffer containing 1 mg/mL Collagenase II, 1 mg/mL Collagenase IV, 0.5 mg/mL Dispase, 10 μg/mL Hoechst, 1 μg/mL ActD, 0.05 mg/mL Proteinase XIV, and 10 U/mL DNase I in perfusion buffer. Cardiomyocyte (CM) and non-cardiomyocyte (non-CM) fractions were separated before fixation. Live CMs were enriched by gravity sedimentation, whereas the supernatant containing non-CMs was transferred to a BSA-blocked 15 mL polypropylene tube. The non-CM fraction was then pelleted at 300 × g for 5 min at 25°C, resuspended in FACS buffer, stained with TO-PRO-3 iodide, and sorted to recover live nucleated single non-CM (Hoechst-positive, TO-PRO-3-negative; Figure S5). The CM pellet was gently resuspended in perfusion buffer with 0.1% BSA using wide-bore pipette tips and kept on ice for short-term handling. When required, cell suspensions were passed through a 100 μm strainer to reduce clumping.

### Cell fixation of adult heart cardiomyocytes and non-cardiomyocytes

After heart cell suspensions were prepared by enzymatic dissociation as described above, cells were fixed using the Parse Biosciences Evercode Cell Fixation Kit according to the manufacturer’s protocol, with the following modifications for adult heart cells. In brief, all polypropylene tubes used during fixation were pre-blocked with 1% BSA for 30 min at room temperature to increase cell retention. For non-CMs, ∼400,000 FACS-sorted cells were pelleted at 200 × g for 10 min at 4°C before fixation. Adult CMs were handled separately from non-CMs and were processed using wide-bore pipette tips. The CM pellet was gently resuspended in 2 mL perfusion buffer containing 0.1% BSA and settled by gravity to enrich viable CMs; this enrichment step was repeated three times before fixation. After permeabilisation, CMs were collected by low-speed centrifugation (30 × g for 3 min at 4°C). Fixed cells were aliquoted and frozen at −80°C in a Mr. Frosty freezing container.

### Pilot mixed-library Parse experiment

In the initial pilot experiment, CM and non-CM fractions prepared from the same heart were combined during fixation. Cells were fixed using Evercode™ Cell Fixation v2 as described above, and libraries were prepared using Evercode™ WT v2. Approximately 48,000 fixed cells/heart were loaded for barcoding, and ∼200,000 barcoded cells were input for library construction, with non-CMs present at roughly two-fold excess over CMs at input, as described in the Results.

### Optimised separated-library Parse experiment

For the optimised workflow, CM and non-CM fractions prepared from the same heart were kept fully separate during fixation and subsequent Parse processing, and were sequenced as independent libraries. Cells were fixed using Evercode™ Cell Fixation v3 as described above. CM libraries were prepared using Evercode™ WT Mini v3, and non-CM libraries were prepared using Evercode™ WT v3. The reason for separating the compartments was to prevent the markedly higher RNA content of CMs from dominating mixed libraries and to permit compartment-specific sequencing-depth allocation. CM and non-CM fractions were loaded within the manufacturer-recommended ranges for the respective kits.

### Sequencing

Libraries were sequenced on a NovaSeq X Plus instrument using paired-end 150 bp reads. The pilot mixed libraries were sequenced with 5% PhiX spike-in, whereas the optimised separated CM and non-CM libraries were sequenced with 10% PhiX spike-in. To test whether limited sequencing depth explained the poor barcode-rank configuration and sequencing saturation of the pilot mixed libraries, one sublibrary was re-sequenced at approximately three-fold greater depth.

### Upstream processing of Parse scRNA-seq data

Upstream processing was performed on AWS using the Parse Biosciences split-pipe workflow with default parameters. The pilot mixed-library data was processed with split-pipe v1.1.2. The optimised separated-library data was processed with split-pipe v1.6.0. For the optimised separated-library data, a GRCm39 reference was generated using split-pipe in mkref mode. Individual sublibraries were processed independently using split-pipe all mode, and outputs from multiple sublibraries were combined using split-pipe comb mode.

### Downstream analysis of Parse scRNA-seq data

Downstream analyses were performed in R 4.5.1 using Seurat 5.4.0. Parse count matrices from the upstream processing were imported with ReadParseBio and converted to Seurat objects with min.cells = 3 and min.features = 40. The mitochondrial transcript fraction was calculated by genes matching the pattern mt-. Per-sample QC distributions of transcript counts, detected genes, and mitochondrial transcript fraction were inspected before filtering. In the optimised separated-library datasets, for the non-CM data, cells were retained if they contained > 900 detected genes, between 500 and 80000 transcript counts, and <15% mitochondrial transcripts. Data were then log-normalised (scale factor 10000), 2000 variable genes were selected, scaled, and PCA was performed. Potential doublets were identified with DoubletFinder using the first 30 PCs. The pK parameter was chosen by parameter sweep to maximise the BCmetric; an expected doublet rate of 2% was assumed. Cells classified as doublets were removed before downstream integration. Filtered singlet objects from individual samples were merged, renormalised, and batch-effect-corrected with Harmony. Neighbor graphs were constructed on the Harmony embedding using PCs 1–30. Resolution 1.2 was selected for the final non-CM analysis. UMAP embedding was computed from the Harmony using PCs 1–30. Non-CM clusters were annotated using canonical marker genes together with FindAllMarkers (min.pct = 0.1, only.pos = TRUE). Significant markers were defined as adjusted p < 0.05. For Figure 4G, marker genes were further restricted to those expressed in at least 50% of cells within the relevant cluster and were ranked by average log_2_ fold change; predicted genes were excluded before plotting. Major annotations included fibroblasts, endothelial cells, macrophages, dendritic cells, pericytes, smooth muscle cells, T cells, B cells, mesothelial cells, Schwann cells, and rare cardiomyocyte contaminants. CM libraries were analysed analogously as a separate compartment, and CM identity was confirmed by expression of *Tnnt2, Myh6*, and *Ryr2*, whereas rare non-CM impurities were identified by markers such as *Cdh5* and *Pecam1*.

### Gene-biotype and rRNA composition analyses

Gene-biotype analyses were performed on the separated CM and non-CM datasets. Gene classes included protein-coding genes, long non-coding RNAs, transcription factors, mitochondrial-encoded genes, nuclear-encoded mitochondrial genes, rRNAs, and pseudogenes. For Figure S2A, a gene was considered detected if it was expressed in at least 1% of cells in the relevant dataset. Because several categories overlap (for example, transcription factors and nuclear-encoded mitochondrial genes are largely protein-coding), percentages do not sum to 100%. For expression-level calculation (Figure S2B), raw counts were aggregated by sample within CMs or non-CMs to generate pseudobulk matrices, normalised to counts per million (CPM), and calculated as the fraction of total CPM of each gene class. For Figures S2C and S2D, rRNA composition was recalculated within the rRNA gene set as CPM_i divided by the sum of CPM across all rRNA genes.

### Comparison with published dataset

Whole-cell CM and non-CM datasets generated in this study were compared with a published mouse left-ventricle snRNA-seq dataset from Li et al.(13). The matrices of control groups were downloaded from GEO (GSE263648), and reanalysed using the similar downstream framework. In pseudobulk comparison, raw counts were aggregated by biological sample within each annotated cell type to generate pseudobulk matrices. Differential expression was performed with DESeq2. For CM comparisons (Figure 5), genes enriched in whole-cell scRNA-seq were defined as log_2_ fold change > 1, adjusted p < 0.001, and baseMean > 1500; genes enriched in snRNA-seq were defined as log_2_ fold change < -1, adjusted p < 0.001, and baseMean > 1500. GO Biological Process enrichment of cardiomyocyte differential-gene sets was performed using g:Profiler, and the top driver terms were visualised. For non-CM comparisons (Figure S3), genes were ranked by -log_10_(p value) × sign(log_2_ fold change), and gene set enrichment analysis (GSEA) was performed with clusterProfiler using GO Biological Process gene sets obtained from msigdbr for Mus musculus. The top positively enriched pathways (normalised enrichment score > 0; enriched in scRNA-seq) and top negatively enriched pathways (normalised enrichment score < 0; enriched in snRNA-seq) were plotted for each cell type.

### Re-analysis of ICELL8 dataset

A published adult mouse heart scRNA-seq dataset generated using ICELL8 platform by Ren et al.(10) was re-analysed to benchmark mitochondrial transcript fractions across cardiac cell populations. The published count matrix was downloaded from GEO (GSE120064) and imported into Seurat. Data were normalised using SCTransform, followed by principal-component analysis. Nearest-neighbor graph construction, clustering, and t-SNE visualisation were performed using PCs 1–20 and a clustering resolution of 1.5. Cell identities were assigned based on canonical marker expression, including cardiomyocytes (*Ryr2, Myh6*), endothelial cells (*Cdh5, Pecam1*), fibroblasts (*Col1a1, Pdgfra*), macrophages (*Mrc1, Fcgr1*), granulocytes (*S100a8, S100a9*), T cells (*Cd3e, Skap1*), B cells (*Ms4a1, Cd79a*), NK cells (*Ncr1, Gzma*), lymphatic endothelial cells (*Lyve1, Prox1*), and epicardial cells (*Wt1, Upk3b*). The mt% values shown in Figure S4 were calculated by genes matching the pattern mt-, and visualised across annotated cell types using violin plots.

### Confocal imaging and volumetric analysis

Freshly isolated CM and non-CM fractions were stained with Pyronin Y to label RNA, MitoViewer to label mitochondria, and Hoechst to label nuclei. Confocal Z-stacks were acquired using a Nikon A1R laser-scanning confocal microscope. 3D volumetric analysis was performed using Imaris software (Bitplane). Surface rendering was applied to the Pyronin Y (RNA), MitoViewer (Mitochondria), and Hoechst (Nucleus) channels to calculate the total volume (µm^3^) of each component per cell.

### Bulk RNA quantification

For bulk RNA analysis, cardiomyocytes and non-cardiomyocytes were sorted and pooled into samples of ∼2×10^5^ cells each. Total RNA was isolated using manufacturer’s TRIzol reagent protocol. RNA concentration was measured using a NanoDrop spectrophotometer, and the average RNA mass per cell (pg) was calculated by dividing the total RNA yield by the input cell count.

## QUANTIFICATION AND STATISTICAL ANALYSIS

Statistical details for individual experiments are provided in the figure legends.

Confocal imaging-based comparisons of cell, nuclear, mitochondrial, and RNA volumes (Figures 2B–2E) and bulk RNA-content comparisons (Figure 2F) were analysed using unpaired, two-tailed Welch’s t tests, as indicated in the figure legends. Data points represent individual cells for imaging experiments and independent mouse replicates for bulk RNA measurements.

For single-cell analyses (Figure 5 and Figure S3), raw counts were aggregated by sample within each cell type to generate pseudobulk matrices, and statistical inference was performed with DESeq2 on those pseudobulk counts. Multiple-testing correction used adjusted p-values as reported by the respective software packages.

For cluster marker identification in Seurat, significant markers were defined as adjusted p < 0.05. For the markers shown in Figure 4G, genes were additionally required to be expressed in at least 50% of cells in the relevant cluster.

## ADDITIONAL RESOURCES

Original adult mouse concomitant CM/non-CM isolation method for tissue dissociation in this study: https://www.ahajournals.org/doi/full/10.1161/CIRCRESAHA.116.309202.

Parse Biosciences user guides for fixation, barcoding, and library construction: https://www.parsebiosciences.com/products/evercode-wt/.

## REFERENCES

1. Ackers-Johnson M, Tan WLW, Foo RSs. Following hearts, one cell at a time: recent applications of single-cell RNA sequencing to the understanding of heart disease. Nat Commun. 2018;9(1):4434.

2. Gardner RS, Tucker NR, Amancherla K. Leveraging Single-Cell Technologies to Advance Understanding of Myocardial Disease. Circ Res. 2026;138(1):e326002.

3. Kim N, Kang H, Jo A, Yoo SA, Lee HO. Perspectives on single-nucleus RNA sequencing in different cell types and tissues. J Pathol Transl Med. 2023;57(1):52–9.

4. Paradis AN, Gay MS, Zhang L. Binucleation of cardiomyocytes: the transition from a proliferative to a terminally differentiated state. Drug Discov Today. 2014;19(5):602–9.

5. Ellman DG, Bjerre FA, Bak ST, Mathiesen SB, Harvald EB, Jensen CH, et al. Protocol to achieve high-resolution single-cell transcriptomics of cardiomyocytes in multiple species. STAR Protoc. 2024;5(3):103194.

6. Gladka MM, Molenaar B, de Ruiter H, van der Elst S, Tsui H, Versteeg D, et al. Single-Cell Sequencing of the Healthy and Diseased Heart Reveals Cytoskeleton-Associated Protein 4 as a New Modulator of Fibroblasts Activation. Circulation. 2018;138(2):166–80.

7. Kannan S, Miyamoto M, Lin BL, Zhu R, Murphy S, Kass DA, et al. Large Particle Fluorescence-Activated Cell Sorting Enables High-Quality Single-Cell RNA Sequencing and Functional Analysis of Adult Cardiomyocytes. Circ Res. 2019;125(5):567–9.

8. Kannan S, Farid M, Lin BL, Miyamoto M, Kwon C. Transcriptomic entropy benchmarks stem cell-derived cardiomyocyte maturation against endogenous tissue at single cell level. PLoS Comput Biol. 2021;17(9):e1009305.

9. Kannan S, Miyamoto M, Zhu R, Lynott M, Guo J, Chen EZ, et al. Trajectory reconstruction identifies dysregulation of perinatal maturation programs in pluripotent stem cell-derived cardiomyocytes. Cell Rep. 2023;42(4):112330.

10. Ren Z, Yu P, Li D, Li Z, Liao Y, Wang Y, et al. Single-Cell Reconstruction of Progression Trajectory Reveals Intervention Principles in Pathological Cardiac Hypertrophy. Circulation. 2020;141(21):1704–19.

11. Nomura S, Satoh M, Fujita T, Higo T, Sumida T, Ko T, et al. Cardiomyocyte gene programs encoding morphological and functional signatures in cardiac hypertrophy and failure. Nat Commun. 2018;9(1):4435.

12. Lu J, Ren J, Liu J, Lu M, Cui Y, Liao Y, et al. High-resolution single-cell transcriptomic survey of cardiomyocytes from patients with hypertrophic cardiomyopathy. Cell Prolif. 2024;57(3):e13557.

13. Li Y, Liu X, Zhang H, Wang Q, Zhu Y, Xu H, et al. Single-nucleus profiling of the left ventricle of the mouse heart after chronic stress. Sci Data. 2025;12(1):1205.

14. Mercer TR, Neph S, Dinger ME, Crawford J, Smith MA, Shearwood AM, et al. The human mitochondrial transcriptome. Cell. 2011;146(4):645–58.

15. Yekelchyk M, Guenther S, Preussner J, Braun T. Mono- and multi-nucleated ventricular cardiomyocytes constitute a transcriptionally homogenous cell population. Basic Res Cardiol. 2019;114(5):36.

16. Loi DSC, Yu L, Wu AR. Effective ribosomal RNA depletion for single-cell total RNA-seq by scDASH. PeerJ. 2021;9:e10717.

17. Pinto AR, Ilinykh A, Ivey MJ, Kuwabara JT, D’Antoni ML, Debuque R, et al. Revisiting Cardiac Cellular Composition. Circ Res. 2016;118(3):400–9.

18. Van Melkebeke L, Verbeek J, Bihary D, Boesch M, Boeckx B, Feio-Azevedo R, et al. Comparison of the single-cell and single-nucleus hepatic myeloid landscape within decompensated cirrhosis patients. Front Immunol. 2024;15:1346520.

19. Gaedcke S, Sinning J, Dittrich-Breiholz O, Haller H, Soerensen-Zender I, Liao CM, et al. Single cell versus single nucleus: transcriptome differences in the murine kidney after ischemia-reperfusion injury. Am J Physiol Renal Physiol. 2022;323(2):F171–f81.

20. Wen F, Tang X, Xu L, Qu H. Comparison of single‐nucleus and single‐cell transcriptomes in hepatocellular carcinoma tissue. Mol Med Rep. 2022;26(5).

21. Nicks AM, Holman SR, Chan AY, Tsang M, Young PE, Humphreys DT, et al. Standardised method for cardiomyocyte isolation and purification from individual murine neonatal, infant, and adult hearts. J Mol Cell Cardiol. 2022;170:47–59.

22. Ackers-Johnson M, Li PY, Holmes AP, O’Brien SM, Pavlovic D, Foo RS. A Simplified, Langendorff-Free Method for Concomitant Isolation of Viable Cardiac Myocytes and Nonmyocytes From the Adult Mouse Heart. Circ Res. 2016;119(8):909–20.

